# Precursor RNA structural patterns at SF3B1 mutation sensitive cryptic 3’ splice sites

**DOI:** 10.1101/2025.02.19.638873

**Authors:** Austin Herbert, Abigail Hatfield, Alexandra Randazza, Valeria Miyamoto, Katie Palmer, Lela Lackey

## Abstract

SF3B1 is a core component of the spliceosome involved in branch point recognition and 3’ splice site selection. SF3B1 mutation is common in myelodysplastic syndrome and other blood disorders. The most common mutation in SF3B1 is K700E, a lysine to glutamic acid change within the pre-mRNA interacting heat repeat domain. A hallmark of SF3B1 mutation is an increased use of cryptic 3’ splice sites; however, the properties distinguishing SF3B1-sensitive splice junctions from other alternatively spliced junctions are unknown. We identify a subset of 192 core splice junctions that are mis-spliced with SF3B1 K700E mutation. We use our core set to test whether SF3B1-sensitive splice sites are different from control cryptic 3’ splice sites via RNA structural accessibility. As a comparison, we define a set of SF3B1-resistant splice junctions with cryptic splice site use that does not change with SF3B1 K700E mutation. We find sequence differences between SF3B1-sensitive and SF3B1-resistant junctions, particularly at the cryptic sites. SF3B1-sensitive cryptic 3’ splice sites are within an extended polypyrimidine tract and have lower splice site strength scores. We develop experimental RNA structure data for 83 SF3B1-sensitive junctions and 39 SF3B1-resistant junctions. We find that the pattern of structural accessibility at the NAG splicing motif in cryptic and canonical 3’ splice sites is similar. In addition, this pattern can be found in both SF3B1-resistant and SF3B1-sensitive junctions. However, SF3B1-sensitive junctions have cryptic splice sites that are less structurally distinct from the canonical splice sites. In addition, SF3B1-sensitive splice junctions are overall more flexible than SF3B1-resistant junctions. Our results suggest that the SF3B1-sensitive splice junctions have unique structure and sequence properties, containing poorly differentiated, weak splice sites that lead to altered 3’ splice site recognition in the presence of SF3B1 mutation.

## Introduction

Splicing is ubiquitous across the human transcriptome. More than 95% of human genes contain one or more intronic sequences that must be removed to produce a fully functional mature RNA (Nilsen and Graveley 2010). These intronic sequences can be processed in multiple ways, through alternative splicing to create multiple isoforms from the same precursor RNA (Pan et al. 2008; Wang et al. 2008). Introns contain several signals that demarcate splice junctions. Core splicing signals include the 5’ splice site, branchpoint, polypyrimidine tract and 3’ splice site (Hastings and Krainer 2001; Black 2003). These precursor RNA signals are recognized during the initial stages of splicing by the spliceosomal U1 and U2 small nuclear ribonucleoproteins (snRNPs) (Wilkinson et al. 2020). In addition, RNA binding proteins (RBPs) can recruit additional splicing factors to guide splice site recognition (Van Nostrand et al. 2020; Tao et al. 2024). Mutations in core spliceosome components and major splicing regulators can dramatically alter splicing. Mutations in SF3B1 and U2AF1, core components of the U2 snRNP, broadly alter splicing and are associated with the development of hematopoietic disorders and cancers (Saez et al. 2017; Zhang et al. 2024). The most common SF3B1 mutation is a change from lysine to glutamic acid at position 700 (K700E) (Wang et al. 2011; Wan and Wu 2013). SF3B1 K700E mutation causes a variety of mis-splicing events, but is specifically associated with cryptic 3’ splicing, a class of alternative splicing where a non-canonical 3’ splice site is recognized. These cryptic 3’ splice sites can be upstream or downstream of the primary canonical 3’ splice site and introduce coding sequence, non-coding sequence and/or premature stop codons (Darman et al. 2015; DeBoever et al. 2015).

All RNAs form co-transcriptional structure based on their primary sequence as they are being transcribed by RNA polymerase (Zamft et al. 2012; Saldi et al. 2021). RNA structure controls access to sequence motifs that regulate interaction with co-factors like RBPs and ribonucleoprotein (RNP) complexes. RNA folding also creates secondary and tertiary structures that can be recognized by RNA interaction partners. Furthermore, RNA folding can also collapse sequentially distant regions into spatial proximity (Zhang and Landick 2016; Hwang et al. 2022; Yu et al. 2023). While RNA structure is clearly important in non-coding RNAs like ribosomal RNA, it has been shown to be an important part of pre-mRNA processing (Novikova et al. 2013; Fabbri et al. 2019). For example, in the *MAPT* precursor RNA, a hairpin at the 5’ splice site spans the U1 interaction motif and is required to control the ratio of exon inclusion and exclusion (Kumar et al. 2022). However, little is known about precursor RNA structures, and because RNA structure is ubiquitous and dynamic, it is difficult to determine which RNA structures are functionally important (Herbert et al. 2023). Here we identify a set of transcripts sensitive to SF3B1 K700E mutation (MT) and characterize them alongside a set of control cryptic 3’ splice sites resistant to SF3B1 mutation (WT) for sequence and structural features that distinguish sensitive and resistant junctions as well as cryptic and canonical splice sites. We find a consistent pattern of structural accessibility at the 3’ splice site for both cryptic and canonical splice junctions. SF3B1-sensitive junctions, particularly the cryptic splice site, are poorly distinguished structurally from the surrounding intron and have weak splice site scores differentiating them from splice junctions unaffected by SF3B1 K700E mutation.

## Results

### SF3B1 K700E mutation consistently causes mis-splicing at a subset of splice junctions

Altering SF3B1 either by knock-down or mutation dramatically changes splicing across the transcriptome (Supp Figure 1A and B). In addition, mis-splicing in SF3B1 K700E mutants is particularly prone to 3’ splice site alteration, resulting in an increase in intron retention and the use of cryptic 3’ splice sites (Figure 1A, Supp Figure 1C). Although SF3B1 strongly perturbs 3’ splice site recognition in multiple different experiments, increased mis-splicing at the 3’ splice site also occurs after perturbation of other U2 snRNP components, such as components of the U2AF complex (Figure 1A, Supp Figure 1C). Supporting the idea that SF3B1 mutants have a dominant negative effect, genes that are mis-spliced in heterozygous SF3B1 K700E datasets are not the same as those affected by knockdown or knockout of SF3B1 (Supp Figure 1D). Characteristically, many SF3B1 mis-splicing events are minor changes (<0.15) to the percent spliced in (PSI) (Figure 1A and B) or are unique to an individual dataset (Figure 1C). In addition, mis-spliced genes verified in patients and proposed to have phenotypic effects are not consistently found in all SF3B1 K700E samples (Supplementary Table 1). Small PSI changes and variability between samples has contributed to difficulty in identifying consistently mis-spliced genes across SF3B1 perturbed samples. However, genes that are consistently mis-spliced by SF3B1 mutation may contain a signature responsible for their sensitivity to SF3B1 K700E. (MDS: GSE63569, K562: GSE187356)

**Figure 1.**
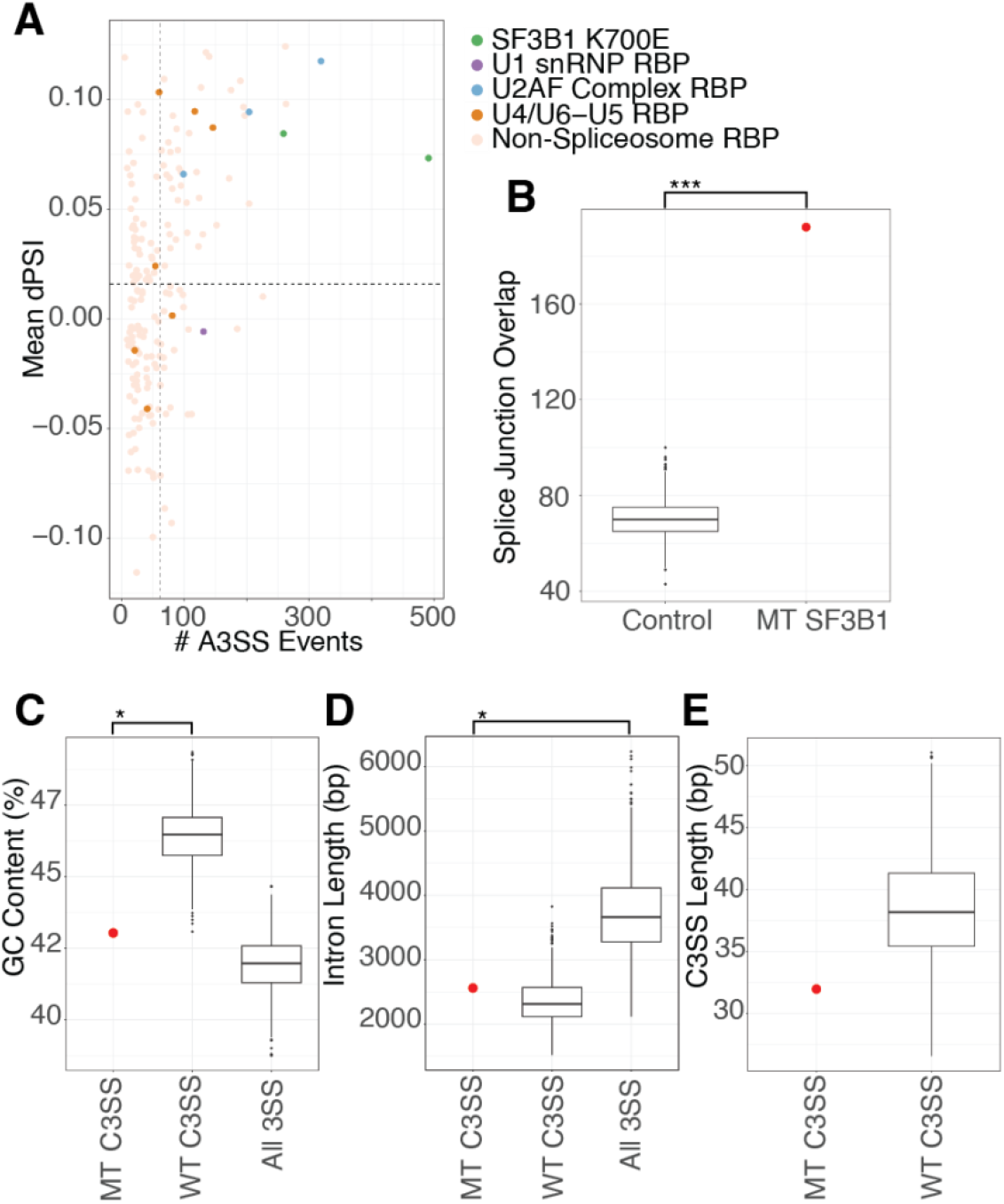
SF3B1 K700E mutation causes consistent mis-splicing at a subset of splice junctions. (A) Degree of alternative 3’ splicing in SF3B1 mutants versus ENCODE RBP knockdown. rMATS analysis from RBP knockdowns in HEPG2 cells were merged and filtered for events with FDR < .09, splice junction counts >10 in knockdown and controls, and deltaPSI > 0.05 and < -0.05. RBPs known to play a large role in splicing are colored with different SF3B1 K700E cell types (green), U1 snRNP components (purple), U2AF complex (blue), U4/U6-U5 tri-snRNP complex (orange) and non-splicing RBPs (tan). (B) Normalized sets of random splice junctions were tested for their overlap and show significantly fewer alternative 3’ splice site events in common than the overlap between sets of splice junctions from SF3B1 K700E datasets. (C) Analysis of GC content in the 150 base pairs upstream of SF3B1 K700E MT-sensitive (MT C3SS) versus WT-resistant cryptic splice sites (WT C3SS). The 192 events shared between MT have significantly lower GC content compared to WT C3SS. WT splice sites selected using random bootstrapping (sampled to 192 events, 1000 times). (D) MT C3SS and WT C3SS have similar intron lengths but are shorter than all other splice junctions. WT C3SS were selected using random bootstrapping. (E) Distance in base pairs between C3SS and paired canonical 3SS in SF3B1 MT sensitive versus resistant C3SS analyzed by bootstrapping. The 192 events shared in SF3B1 MT show no significant difference in C3SS length compared to WT C3SS (sampled to 192 events, 1000 times). (*** = p < .0005, **= p < .005, *= p < .05)

To identify a subset of genes that are consistently altered by SF3B1 K700E, we sequenced pre-B NALM-6 cell lines isogenic for SF3B1 WT and SF3B K700E. Our sequencing data shows high similarity to gene expression and splicing patterns from previous sequencing of NALM-6 SF3B1 K700E lines (Supp Figure 1G) (Darman et al. 2015). We analyzed our NALM-6 sequencing data in combination with RNA-sequencing from published studies in the pre-B K562 cell lines with normal SF3B1 and SF3B1 K700E. For further validation, we compared our cell-line splicing results with data from a set of patients with myelodysplastic syndrome (MDS), an early B-cell clonal deficiency. Using cell-line and MDS patient datasets allows us to capture natural splicing variation while minimizing the impact of genetic variation in patients and the noise of differential gene expression from different tissues. Due to the disproportionate increase in 3’ cryptic splice site usage with SF3B1 mutation, we concentrate on splice junctions containing a cryptic 3’ splice site for further analysis. We find 192 cryptic 3’ splice sites in 192 genes to be mis-spliced in two or more SF3B1 K700E datasets with a dPSI > .05 using rMATS analysis (Supplementary Table 2). This overlap in number of alternatively spliced genes between three different experiments is significantly more than would be expected by chance (Figure 1B). Both LeafCutter and rMATs splicing analysis produced similar results (Supp Figure 1E). We identified several previously published junctions in our high confidence set, including BCL2L1, MAP3K7, and DVL2 (Zhao et al. 2021; Fuentes-Fayos et al. 2022; Lieu et al. 2022). Kegg analysis reveals a significant enrichment of these genes in the nucleocytoplasmic transport pathway (9.9-fold enrichment, Supplementary Table 3) including IPO7, TNPO3 and TPR (Golomb et al. 2012; Kosar et al. 2021; Iwanami et al. 2023). Interestingly, gene ontology analysis of biological processes reveals a significant enrichment in mRNA transport (6.7-fold) and histone modification (3.2-fold, Supplementary Table 4) (Ge et al. 2020). Disruption of SF3B1 results in perturbation of RNA processing, similar to the impact of disrupting many other RNA binding proteins, but for SF3B1, this perturbation may contribute to the diversity of RNA processing in individuals with SF3B1 mutations (Hentze et al. 2018).

### Sequence patterns in SF3B1-sensitive splice junctions

SF3B1 K700E spliceosomes are not known to have strong alternative sequence preferences at the branch-point or 3’ splice sites (Gupta 2019). To determine whether SF3B1-sensitive MT splice junctions have unique sequence properties, we compared them to control splice junctions containing cryptic 3’ splice sites with similar PSI properties that are SF3B1-resistant (WT junctions). We identified 2801 SF3B1-resistant WT junctions with cryptic splice sites. We found that MT splice junctions and WT splice junctions have significantly different GC content around the 3’ splice site (Figure 1C). Both MT and WT junctions have shorter introns than junctions without alternative 3’ splice sites (Figure 1D). When we measure the distance between the cryptic 3’ splice site to the canonical 3’ splice site, we find that SF3B1-sensitive junctions are not statistically significantly different from WT junctions (Figure 1E). Based on the difference in GC content between MT and WT splice junctions, we decided to further analyze the splice site strength and nucleotide composition at the cryptic and canonical sites of these junctions.

We analyzed the strength of each splice site using the 3’ MAXENT score and found that the WT and MT junctions were significantly different (Figure 2A). This is consistent with our GC content analysis but highlights the fact that while the WT canonical site is much stronger than the MT canonical site, both the WT and MT cryptic splice sites are weak (Figure 2A). The MAXENT score for the 3’ splice site includes the last 20 bases of the intron and first three bases of the exonic sequence (Yeo and Burge 2004). To determine what was driving the differences between the WT and MT scores, we analyzed the nucleotide composition around both the cryptic and canonical splice sites. We found that the canonical splice sites of the MT and WT junctions were similar (Supp Figure 2A, Figure 2D & E). In contrast, the MT and WT cryptic splice sites differed in sequence content (Figure 2B and C). The MT cryptic splice site dinucleotide is embedded within an extended polypyrimidine tract that flanks both sides of the junction (Figure 2B), while the WT cryptic site follows a more typical polypyrimidine tract pattern that is U/C rich upstream of the splice site (Figure 2C). Further analysis of nucleotide composition supports sequence differences in the proportion of adenines (A) upstream of the MT and WT cryptic splice sites (Figure 2D, top) but no differences in adenines between MT and WT canonical splice sites (Figure 2D bottom). Likewise, the percentage of uridine (U) is decreased upstream of the cryptic splice site and increased downstream of the cryptic splice site between the MT and WT junctions (Figure 2E, top), but is similar between MT and WT canonical splice sites. These results suggest that the difference between the MT and WT splice scores reflects nucleotide composition outside of the immediate area around the canonical sites (Figure 2D and E, bottom). Although we found a difference in overall GC content, we did not detect a difference in the pattern of G or C nucleotides in the immediate area of the cryptic or canonical splice site dinucleotides (Supp Figure 2A). Our analysis of the sequence features of SF3B1 sensitive junctions supports the hypothesis that genes that are alternatively spliced by SF3B1 mutants have unique sequence properties.

**Figure 2.**
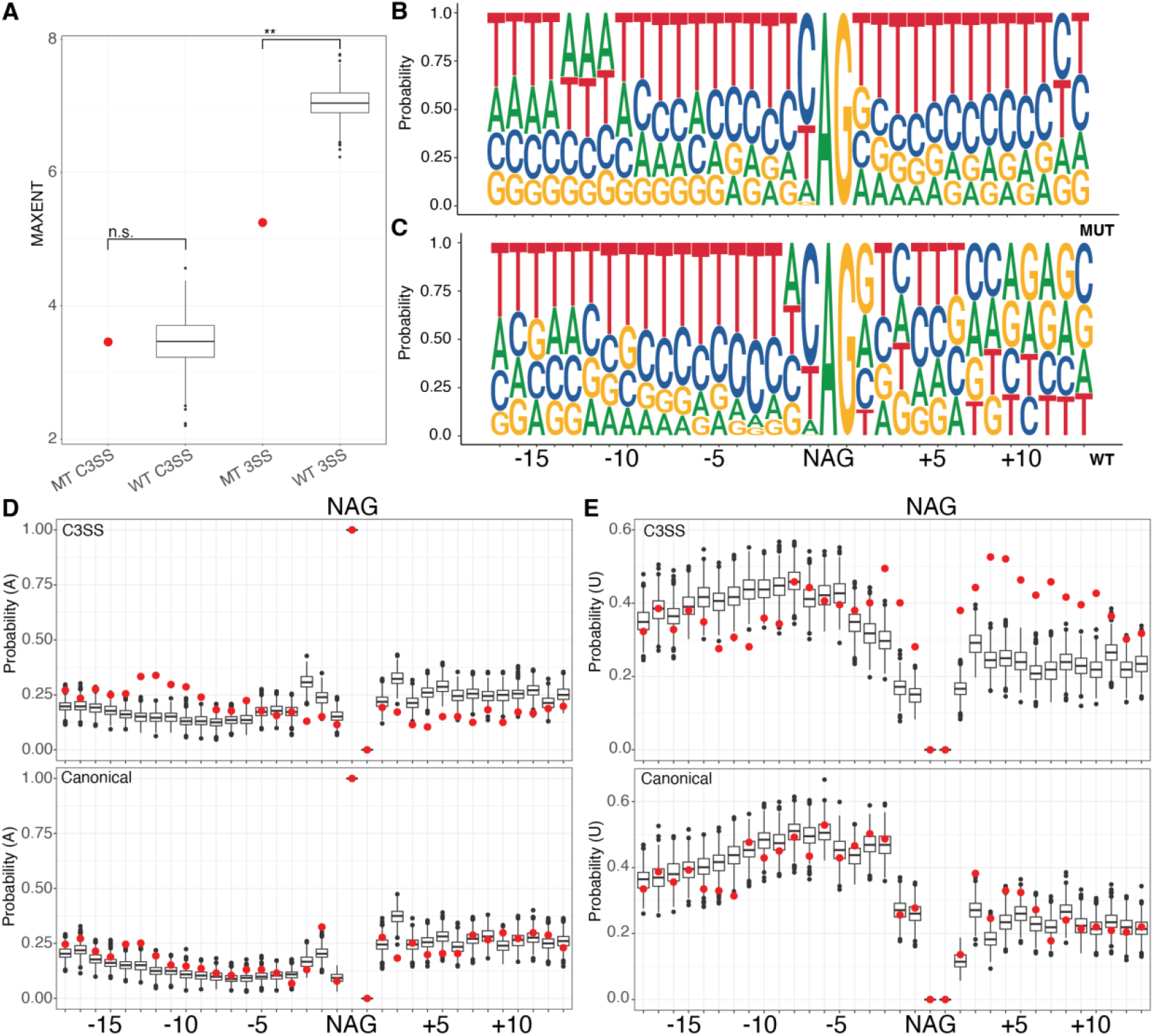
SF3B1 sensitive MT cryptic splice sites are embedded in an extended polypyrimidine tract. (A) Splice site strength was calculated using MAXENT at cryptic splice sites (C3SS), with both MT and WT having similar low strengths at the cryptic position. In contrast, the same measurement at canonical splice sites (3SS) is significantly higher at the WT splice site than at MT splice sites. WT splice sites were selected using random bootstrapping. (B) The MT cryptic splice sites (top) are embedded in an extended polypyrimidine tract compared to paired canonical splice sites (bottom) as shown by sequence logo analysis of the nucleotide probability. (D) Probability of adenines and (E) uridines flanking MT C3SS (red, top) and paired 3SS (red, bottom) compared to WT C3SS (black, top) and 3SS (black, bottom) indicating changes in nucleotide composition, particularly an increase in uridines downstream from the splice site motif (NAG) at the MT C3SS. WT splice sites were selected using random bootstrapping. (*** = p < .0005, **= p < .005, *= p < .05)

### Conserved accessibility patterns at the 3’ splice site motif

RNA structure influences regulation of pre-RNA processing (Herbert et al. 2023). We performed chemical probing to determine the *in vitro* structural accessibility for 83 MT splice junctions, 38 WT splice junctions and 15 control splice junctions with no known cryptic 3’ splice sites. We selected these 83 junctions from the core set of 192 SF3B1-sensitive junctions, but not all junctions could be synthesized due to complexity. Selected junctions had the same properties as the larger subset used to analyze nucleotide composition (Supp Figure 1F). Replicates of chemical probing experiments resulted in highly correlated reactivity profiles (Supp Figure 3A). To determine whether *in vitro* precursor RNA structure can be used as a proxy for *in vivo* RNA structure, we isolated nuclei and performed chemical probing to develop reactivity profiles across the cryptic and canonical splice sites for five of our splice junctions. All RNAs had high correlation of reactivity data between *in vivo* and *in vitro* conditions (Supp Figure 3A). Thus, we used our *in vitro* data to analyze RNA structure at the cryptic and canonical splice sites in both WT and MT splice junctions. When we quantified the reactivity around the NAG splicing motif, we found a consistent pattern where the N is significantly less reactive than average across the region, the adenine is highly reactive, and the guanine is average (Figure 3A, yellow highlight). The same pattern is visible for the cryptic and canonical splice sites in both MT (Figure 3B, purple and orange) and WT splice junctions (Figure 3C, purple and orange) with the N significantly less reactive than the adenine.

**Figure 3.**
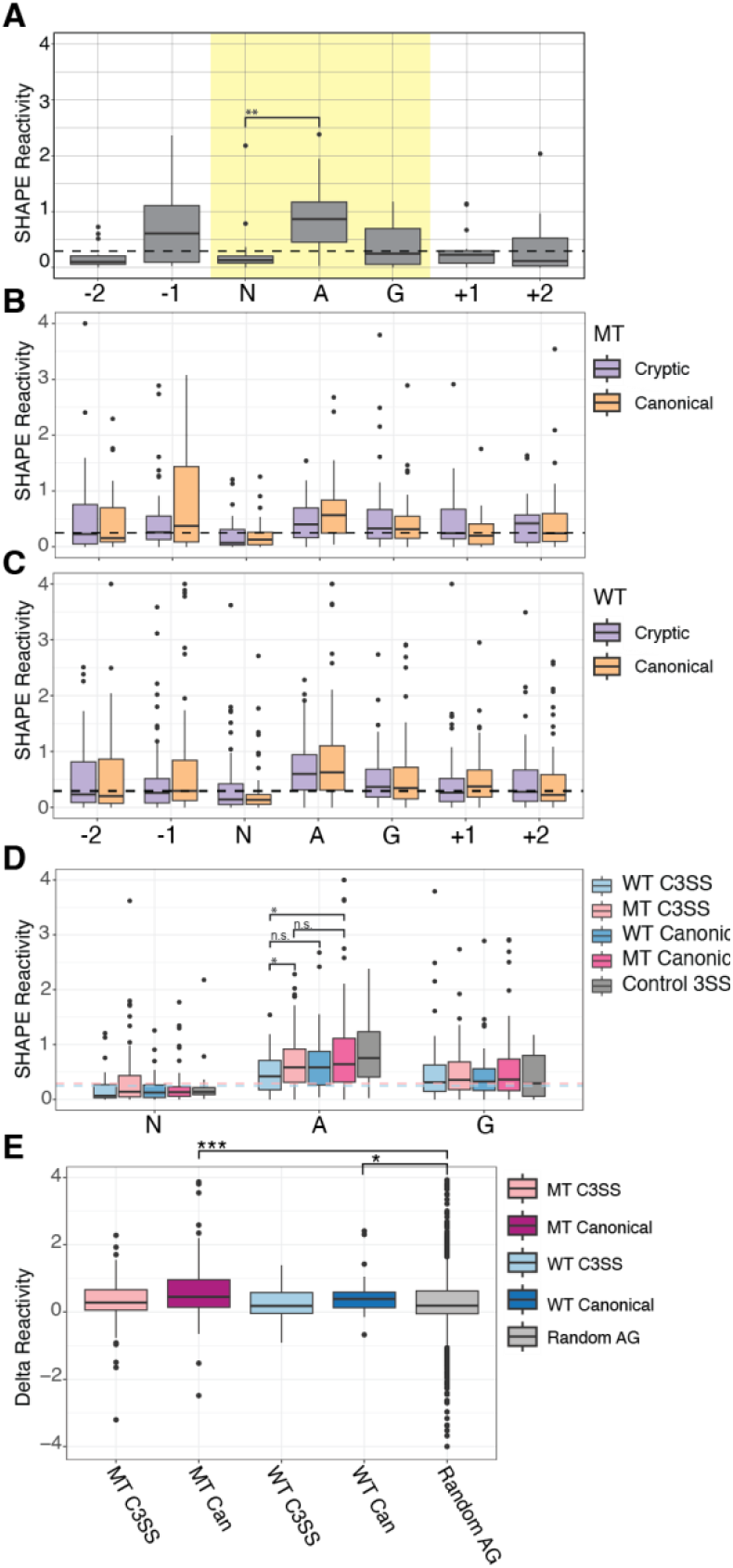
Conserved pattern of accessibility at cryptic and canonical 3’ splice sites. (A) Control 3’ splice sites (3SS) with no alternative splicing show a characteristic accessibility pattern from chemical probing SHAPE reactivity data with the central adenine in the NAG motif significantly higher than the preceding N. The median reactivity from ±150 surrounding bases is plotted as a reference (black) (n=15). (B) SHAPE reactivity profile of SF3B1 MT sensitive C3SS plotted next to paired canonical SHAPE reactivities and the median reactivity from ±150 splice site bases plotted as a dashed black line (n = 83). (C) SHAPE reactivities of WT C3SS and paired canonical splice sites and the median reactivity from ±150 splice site bases plotted as a dashed black line (n = 39). (D) SHAPE reactivities from A, B, and C plotted together, significance tested with a pairwise Wilcoxon test, and the median SHAPE reactivities for ±150 splice site bases from MT C3SS (pink) and WT C3SS (blue) plotted as dashed lines. (E) SHAPE reactivities for -120 to +15 from 3SS normalized to control splice junctions. (*** = p < .0005, **= p < .005, *= p < .05)

Prior computational predictions have suggested that the guanines within cryptic MT junctions are less accessible than canonical MT junctions (Kesarwani et al. 2017). Reactivity data from chemical probing experiments directly measures nucleotide accessibility and can be compared to base-pairing probability predictions (Deigan et al. 2009). We found no significant difference in reactivity at the guanine in cryptic and canonical splice sites in either the MT or WT splice junctions (Figure 3D). However, the central adenine of the MT cryptic site is significantly less reactive than the central adenine of the WT cryptic site (Figure 3D). The MT cryptic site adenine is also less reactive than the MT canonical site (Figure 3D). The same trend is present for WT cryptic site adenine compared to its more accessible partner canonical site (Figure 3D) although this is not significant. In addition, at the NAG splicing motif, we find that the magnitude of difference between the less reactive N and the highly reactive A is lower for cryptic splice sites than for canonical splice sites, with canonical sites being significantly more reactive than non-splice site control NAGs (Figure 3E). While we find the adenine to be more relevant in our experimental data, the principle of a less accessible cryptic splice site is maintained in our results, similar to the prior study (Kesarwani et al. 2017).

We normalized our reactivity data to other NAG sequences outside of the annotated splice sites (Figure 4A and B). Non-splice site NAG motifs generally have reactive central adenines, but MT canonical splice sites remain more reactive while the preceding N in the NAG splicing motif is less reactive (Figure 4A). The same pattern is present in the canonical WT splice site although the central adenine is not as reactive (Figure 4B). To further study RNA structure at the 3’ splice site, we incorporated our MT and WT experimental data into computational models of co-transcriptional and post-transcriptional folding. Experimental structural reactivity can significantly improve computational predictions (Siegfried et al. 2014). However, we found that computational co-transcriptional models that fold short sequences (66 nts) can contain artifacts where the 3’ end tends to be more accessible regardless of the sequence (Supp Figure 3B). Folding longer windows of sequence around the 3’ splice site with strict pairing to maintain a model of co-transcriptional folding eliminates this positional bias (Supp Figure 3C). Incorporating reactivity data into a position independent co-transcriptional model, we find cryptic 3’ splice sites trend toward less accessibility than canonical 3’ splice sites (Figure 4C, Supp Figure 3D). Post-transcriptional models follow the same trend with the central adenine of the cryptic splice site less accessible than the canonical splice site in both WT and MT junctions (Supp Figure 3E). Interestingly, other non-splice site NAG sequences within our data share similar high reactivity at the adenine (Figure 4C and D). The pattern of reactivity data, showing inaccessible Ns and reactive As at splice sites, is not reproduced in the base-pairing probabilities after computational modeling, even with experimental data incorporated, suggesting that the sequence context of cryptic and canonical is influencing structure, particularly in the co-transcriptional model with tight distance constraints (Figure 4C and D, Supp Figure 3D). Overall, like in reactivity data, canonical splice sites trend toward higher accessibility than cryptic splice sites (Supp Fig 4A).

**Figure 4.**
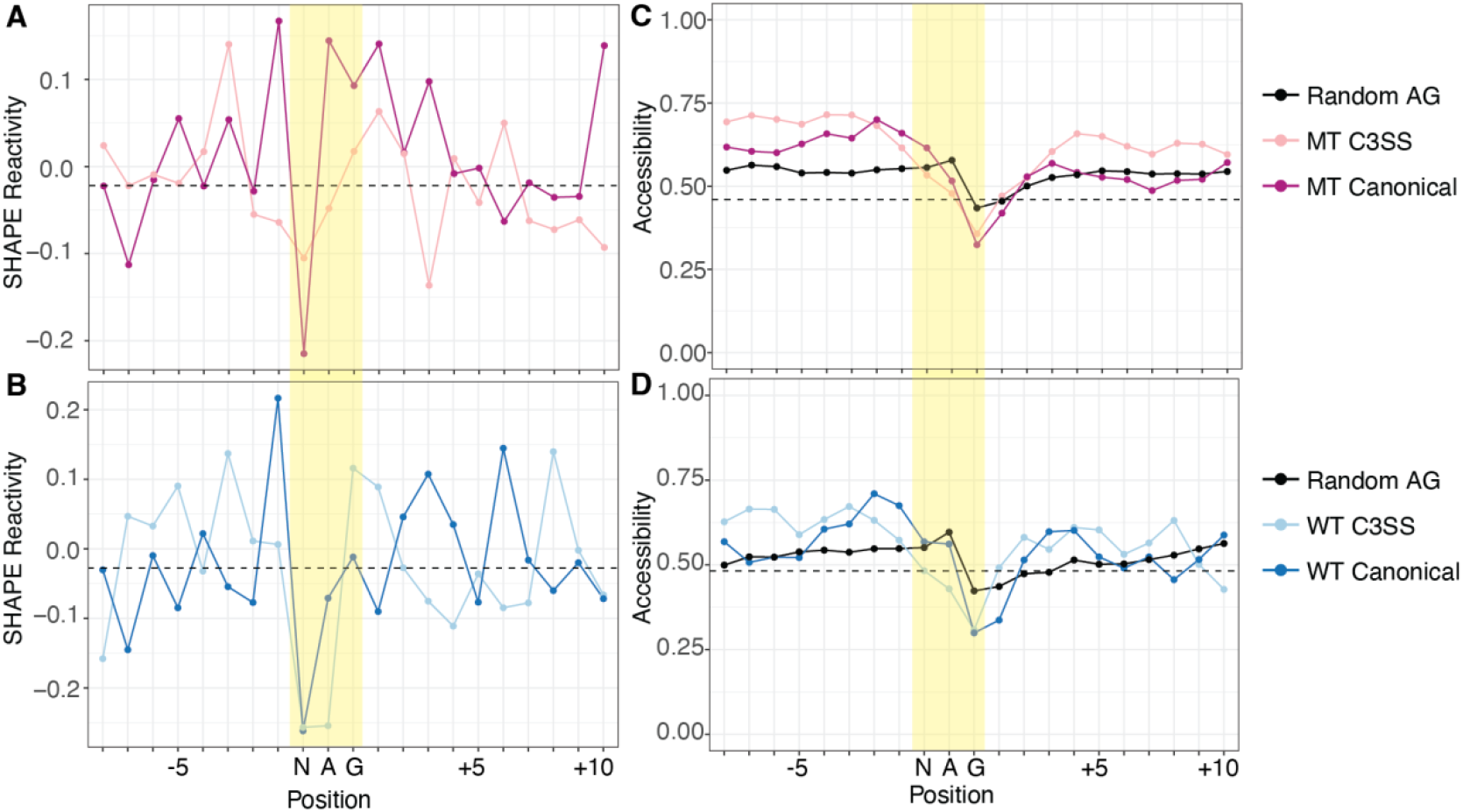
Cryptic and canonical splice sites have similar SHAPE reactivity and predicted accessibility. (A) Dip in SHAPE reactivity at the N of the NAG splice site motif for both MT C3SS (light pink), paired canonical (dark pink) and (B) WT C3SS (light blue) and paired canonical (dark blue) compared to flanking nucleotides. Data normalized to mean SHAPE reactivity of random intronic or exon non-splice site AG motifs. Median normalized reactivities for ±10 bases from the MT and WT cryptic splice site plotted as black dashed horizontal lines. (C) Base accessibility exhibits a downward trend over the NAG splice site motif compared to flanking nucleotides for both MT C3SS (light pink), paired canonical (dark pink) and (D) WT C3SS (light blue) and paired canonical (dark blue). Base accessibility was determined through SHAPE-guided RNAfold predictions averaged over max distances of 27 to 34.

Accessibility at the NAG splice site is limited to only three nucleotides and may not represent the accessibility of the overall junction. In fact, we find that MT cryptic 3’ splice sites upstream of the NAG motif are significantly more accessible than random NAG motifs (Supp Figure 4A). To further investigate the surrounding region, we incorporated experimental reactivity data into scanFold to analyze intronic sequences around the splice sites for stable structures and ensemble diversity (Andrews et al. 2018). We found that MT splice junctions contain significantly less stable structures across both cryptic or canonical splice sites than their partner sites in WT splice junctions (Figure 5A and C). Supporting this observation, MT splice junctions have significantly higher ensemble diversity at both cryptic and canonical splice sites than their partner WT splice sites (Figure 5B and C). Within WT splice junctions that are resistant to splicing changes in the context of SFB31 mutation, cryptic sites are significantly more stable, although this difference is minimal and does not correspond to a significant change in ensemble diversity (Figure 5C and D). To corroborate this, we analyzed the amount of base-pairing in co-transcriptional minimum free energy structures surrounding the NAG motif. We find that there is significantly less base-pairing around MT junctions than in WT or control junctions at both the cryptic and canonical sites (Figure 5E). In contrast to the scanFold calculations, WT canonical splice sites have more base-pairing than WT cryptic splice sites (Figure 5E). Overall, this structural data suggests that cryptic NAG motifs are less accessible than NAG motifs in their canonical counterparts (Figure 4E), but more robust splice junctions, typified by cryptic and canonical WT SF3B1 resistant regions, have more stable structure, less ensemble diversity and more base-pairing outside of the immediate NAG splicing motif (Figure 5C-E).

**Figure 5.**
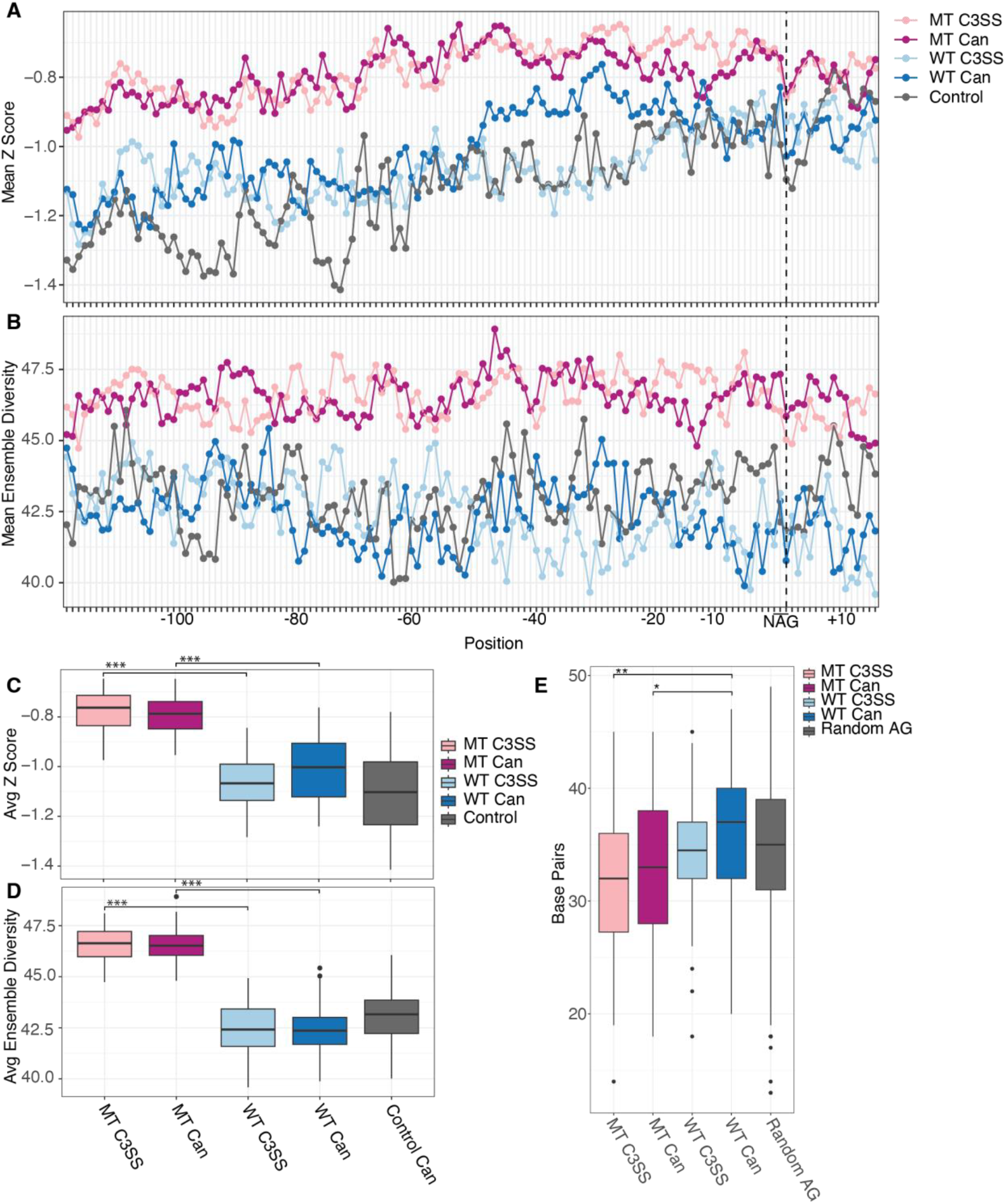
Increased flexibility of upstream intronic sequences in SF3B1 MT sensitive 3’ splice sites. (A) Higher Z scores suggest the presence of less stable structures upstream of NAG splice site motifs for MT cryptic 3’ splice sites (MT C3SS, light pink) and paired canonical (MT Can, dark pink upstream of NAG splice site motifs compared to WT C3SS (light blue) and WT Canonical (dark blue). (B) Likewise, higher ensemble diversity indicates the propensity of MT C3SS (light pink) and MT canonical splice sites to form multiple distinct secondary structures compared to WT C3SS (light blue) and paired WT canonical splice sites (dark blue). Control 3SS (gray). (C) Significantly less stable MT regions (light pink, dark pink) compared to WT C3SS (light blue) and paired canonical (dark blue). Box plot of Z scores averaged per base for bases -121 to +15 across all MT C3SS, paired MT canonical, WT C3SS, paired WT canonical and control 3SS (gray). (D) Significant difference in ensemble diversity between SF3B1 MT C3SS (light pink) paired MT canonical (dark pink), WT C3SS (light blue) and WT canonical (dark blue). Ensemble diversity plotted as described in (C). (E) Significant difference in the number of paired bases in MFE structures of MT C3SS (light pink) and MT canonical (dark pink) compared to WT C3SS (light blue) and WT canonical (dark blue). The number of paired bases for -121 and +15 bases flanking cryptic and canonical splice sites was determined from SHAPE guided MFE structures produced with RNAfold. (*** = p < .0005, **= p < .005, *= p < .05)

## Discussion

Here we describe the structural pattern of more than 125 splice junctions, the majority of which contain a canonical and a cryptic 3’ splice site. This is the largest set of RNA structure data at 3’ splice junctions and is highly correlated to our *in vivo* validation subset. Analyzing our reactivity data, we find that 3’ splice junctions have a characteristic pattern of accessibility at the NAG splicing motif where central adenine is more reactive than the preceding N. This pattern is present in both cryptic and canonical sites and also in both MT and WT splice junctions. However, we find that cryptic splice sites have lower magnitude changes in reactivity between the N and the A than non-splice site NAGs. We propose that the NAG pattern of reactivity is necessary to demarcate a strong splice site and that the magnitude of change between the N and the A may distinguish the splice site from the surrounding intron. Computational modeling of introns, even with reactivity data, does not capture the experimental reactivity pattern at the splice site. The transition between low and high reactivity at the N and A nucleotides in the experimental data is abrupt and may be due to lack of flexibility at the nucleotide rather than direct base-pairing and slight differences in nucleotide reactivity (Wilkinson et al. 2009). This could account for why the reactivity data is not reflected in the base-pairing probability from computational modeling. However, we find that the pattern of accessibility using computational modeling is similar for cryptic and canonical splice sites in both MT and WT splice junctions, like in our reactivity data. Derivation of experimental reactivity data on intronic sequences is important for direct analysis of nucleotide accessibility and to best inform computational modeling.

In addition to the structural characteristics at the NAG splice site, we also find that MT splice junctions are generally more flexible with higher ensemble diversity, lower structural stability and less base-pairing than WT or control splice junctions. This increase in flexibility covers a large area around the splice junction. While an increase in RNA structure in the WT splice junctions may seem counterintuitive for promoting splicing, there are other examples where RNA structure forms and promotes accessibility of specific regions. For example, *MBNL1* mRNA is bound by its own product, MBLN1 protein, which causes downstream restructuring of a functional RNA element (Bubenik et al. 2020). Likewise, in the 5’ UTR of *SERPINA1* mRNA, structures upstream of the start codon are necessary to promote accessibility of the Kozak sequence (Grayeski et al. 2022). Less flexibility in WT splice junctions may promote splicing or recruit specific RNA binding proteins during early splice site recognition compared to MT splice junctions.

SF3B1 mutation is common in myelodysplastic syndrome and other blood disorders, but it is not clear how this confers an advantage on mutant cells. Previous studies have looked for a role of specific mis-spliced genes in oncogenic phenotypes or proposed roles other than splicing in SF3B1 disruption, such as R-loop pathways (Jiang et al. 2023). We analyzed whether MT genes that are consistently mis-spliced by SF3B1 K700E have similar structural characteristics to further understand the mechanism underlying the cohort of mis-spliced genes. We find that MT splice junctions have weak splice site strength at both the cryptic and canonical splice sites. This corresponds with a lack of differentiation in reactivity between the splice site NAG nucleotides for MT splice junctions, particularly the cryptic splice site compared to non-splice site NAG motifs. In addition, we find that MT splice junctions are significantly more flexible than WT splice junctions. Our results show that multiple features differentiate MT splice junctions from WT splice junctions, suggesting that recognition of splice junctions is likely due to multiple factors across a broad region of the precursor mRNA encompassing both the cryptic and canonical splice sites.

## Acknowledgments

We thank the Genomics and Bioinformatics Core of the Clemson University Center for Human Genetics for analytical and computational assistance with Secretariat High Performance computing cluster (supported by NIH grant P20GM139769) used for data analysis.

## Materials and Methods

### Cell culture

Human Nalm-6 SF3B1 K700E mutant and wildtype cells are from Horizon Discovery (Lafayette, Colorado). Nalm-6 cells were maintained in RPMI-1640 medium (Gibco) supplemented with 10% fetal bovine serum (FBS), 1% penicillin-streptomycin at 37 C with 5% CO2 and passaged every three days.

### Library preparation and RNA sequencing

Whole cell RNA was extracted from 1×10^6^ SF3B1 K700E and wildtype cells using standard Trizol extraction followed by DNA digestion and rRNA depletion prior to reverse transcription (ThermoFisher TRIzol, 5PRIME PhaseLock Heavy, Invitrogen Purelink RNA columns, ThermoFisher Scientific TurboDNAse Kit, and Qiagen FastSelect -rRNA depletion kit). Briefly, 1 ug of RNA was incubated with QIAseq FastSelect -rRNA HMR in two-minute steps decreasing in 5 C increments from 75 to 55 C and finishing with 37 C for two minutes followed by 25 C for two minutes. This reaction was put straight into RT. Reverse transcription was performed using SuperScript IV reverse transcription kit with random nonomers, and second strand synthesis was completed with the Second Strand cDNA Synthesis Kit (Invitrogen, ThermoFisher Scientific). cDNA was purified with the Monarch PCR and DNA cleanup kit (New England Biolabs, NEB) and sequencing libraries were prepared under standard protocol using the Nextera DNA Library Preparation Kit (Illumina). RNA purification and cDNA libraries were prepared in triplicate for Nalm-6 SF3B1 mutant and wildtype cells grown on different days. Paired-end libraries (2 × 150) were sequenced on an Illumina NovaSeq.

### Splice junction characterization and quantification

Specific parameters and scripts are available on GitHub (herber4/SF3B1_C3SS_Structure_Paper.git). RNA sequencing reads from MDS patient samples (MDS: GSE63569, SF3B1 K700E n=8, SF3B1 WT n=4, CD34+ healthy n = 5) and K562 SF3B1 K700E isogenic cell line (GSE187356, SF3B1 K700E n=2, SF3B1 WT n=2) were downloaded from the Sequence Read Archive (Dolatshad et al. 2015; Lieu et al. 2022) and our NALM sequencing data all underwent the same analysis pipeline. Bbduk from bbmap was used to trim reads of adapters and to discard low quality reads and any remaining rRNA reads. Quality trimming was performed using bbduk (BBMap v38.73) using parameters provided at (bbduk_trim.sh) (Bushnell 2014). Reads were mapped to Hg38 using Hisat2 v2.1.0 under default parameters and alignment files were sorted and indexed using Samtools v1.10 (hisat_alignment.sh) (Li et al. 2009; Kim et al. 2019). Splicing analysis was conducted using rMATs v4.1.1 on the SF3B1 K700E samples against their matched controls with parameters allowing for novel splice site detection and variable read length (rmats.sh) (Wang et al. 2024). Alternative 3’ splicing events shared in two or more cell types with and FDR < .1 and deltaPSI > .05 were included for further analysis. We identified resistant cryptic 3’ splice sites without splicing changes in SF3B1 mutant cells. First, junctions were extracted from mapped bam files using the regtools junctions extract command allowing for a minimum anchor length of 8 base pairs, and a minimum and maximum intron size of 50 and 500000, respectively (https://github.com/griffithlab/regtools.git). Then, junctions identified in the previous step were annotated with the Gencode Hg38 v39 gtf file using the regtools junctions annotate command under default parameters. Finally, identification and filtering of cryptic 3’ splice sites in SF3B1 WT cells was conducted in R, resulting in the identification of 2801 C3SS events. These 2801 events were used as controls for all further analyses of sequence characteristics. Events were filtered and compared to previously published ENCODE RBP knockdown rMATS (https://github.com/flemingtonlab/SpliceTools.git) (Flemington et al. 2023). GC content, intron length, C3SS length, sequence content and bootstrapping analyses were all conducted in R v4.4.2. Maximum entropy scores for cryptic and paired canonical splice sites were generated with MAXENT and further analyzed for significance in R (Yeo and Burge 2004).

### gBlock Design, in-vitro & in-vivo SHAPE Mapping

Gene fragments were designed and synthesized by Integrated DNA technologies (IDT). Each gene fragment contains 450 base pairs flanking both sides of the 3’ splice site, with a T7 promoter on the 5’ end and both ends capped with stabilizing hairpin sequences. Using the gene fragments as template, in vitro RNA synthesis was performed for two hours at 37 C using the HiScribe T7 High Yield RNA Synthesis Kit (New England Biolabs). In vitro synthesized RNA was DNA digested with TurboDNAse and purified with RNA XP cleanup beads (Beckman Coulter). RNA was incubated in folding buffer (400 mM bicine, 200 mM NaCl, 20 mM MgCL2) and 25 mM 5NIA or DMSO at 37 C for 5 minutes and immediately cleaned up using RNA XP beads (Beckman Coulter) and input into error-prone reverse transcription using the Invitrogen Super Script II (ThermoFisher Scientific) with modified RT buffer (10X: 500mM Tris pH 8, 750 mM KCl) supplemented with 6 nM manganese, RNAse inhibitor, and reverse primer 5-TGTTGGAGTCACTCGACTCCGGT-3. The RT product was cleaned up with the RNA XP beads and used as template for the Second Strand cDNA Synthesis kit (Invitrogen, ThermoFisher Scientific). Finally, libraries were synthesized with standard Nextera DNA library prep and sequenced (2 × 150, Illumina).

### In vivo validation of RNA structure

Nalm-6 SF3B1 K700E mutant cells were grown to confluency, 1 × 10^7^ cells were pelleted, and nuclei were fractionated using the Cytoplasmic and Nuclear RNA purification kit (Norgen Biotek). Briefly, cell pellets were treated with lysis buffer J on ice for 15 minutes, and nuclei separated by centrifugation at 600 g for 5 minutes. Remaining nuclei were resuspended in 1X buffer SK then treated with 1X folding buffer, 200 U murine RNAse inhibitor (NEB) and either 25 mM 5NIA or DMSO and incubated at 37 C for 10 minutes. Finally, RNA was purified with the remaining steps of the Norgen purification kit and DNAse digested as previously described. Error-prone RT was conducted as previously described with the addition of random nonomers instead of gene specific primers, and RT products were cleaned up using the RNA XP beads. To amplify target preRNA, subsequent PCR reactions using gene specific primers were used to incorporate Illumina barcodes (Supplementary Table 5). Both PCR reactions were conducted for 15 cycles each with the Monarch Q5 High-Fidelity DNA Polymerase (NEB). Final PCR products were pooled and size selected from a DNA agarose gel with the Monarch DNA Gel Extraction Kit (NEB). Libraries were sequenced at 2 × 150 on an Illumina MiniSeq. SHAPE profiles were generated with shapemapper2 v2.2.0(29114018) and technical replicates with high correlation were merged for further analysis. Correlation analysis between in vivo and in vitro SHAPE profiles was conducted in R.

### SHAPE analysis

SHAPE analysis was conducted with shapemapper2 v2.2.0 under default parameters using DMSO controls as untreated and 5NIA reads as modified, respectively (shapemapper.sh, 29114018). Normalized SHAPE reactivity profiles were adjusted to between 0 and 4 and utilized for all further downstream analyses. SHAPE guided base pairing probabilities for max distances of 27 through 34 were predicted using the RNAfold function of ViennaRNA v2.4.18 and the mean of this window was used for further analysis. Structural trends flanking the cryptic and canonical splice sites were assessed by predicting SHAPE guided MFE structures for the last 120 bases of the intron and the first 15 bases of the exon using RNAFold function with no max distance (22115189, 38499485). Dot bracket files produced in the previous step were imported into R, and hairpin length was determined as the sum of paired bases divided by two. Structural trends in this region were further assessed by evaluating Z scores and ensemble diversities predicted with Scanfold (PMID: 31711930). Scanfold was run under default parameters using SHAPE data to guide predictions and allowing a window size of 200 and step size of 20. Windows from both cryptic, canonical and control splice sites were analyzed. Per base z scores and ensemble diversities were averaged over all splice sites included in each group and visualized in R. Significance between nucleotide reactivity was assessed with Wilcoxon test.

---

**Supplementary Figure 1.**
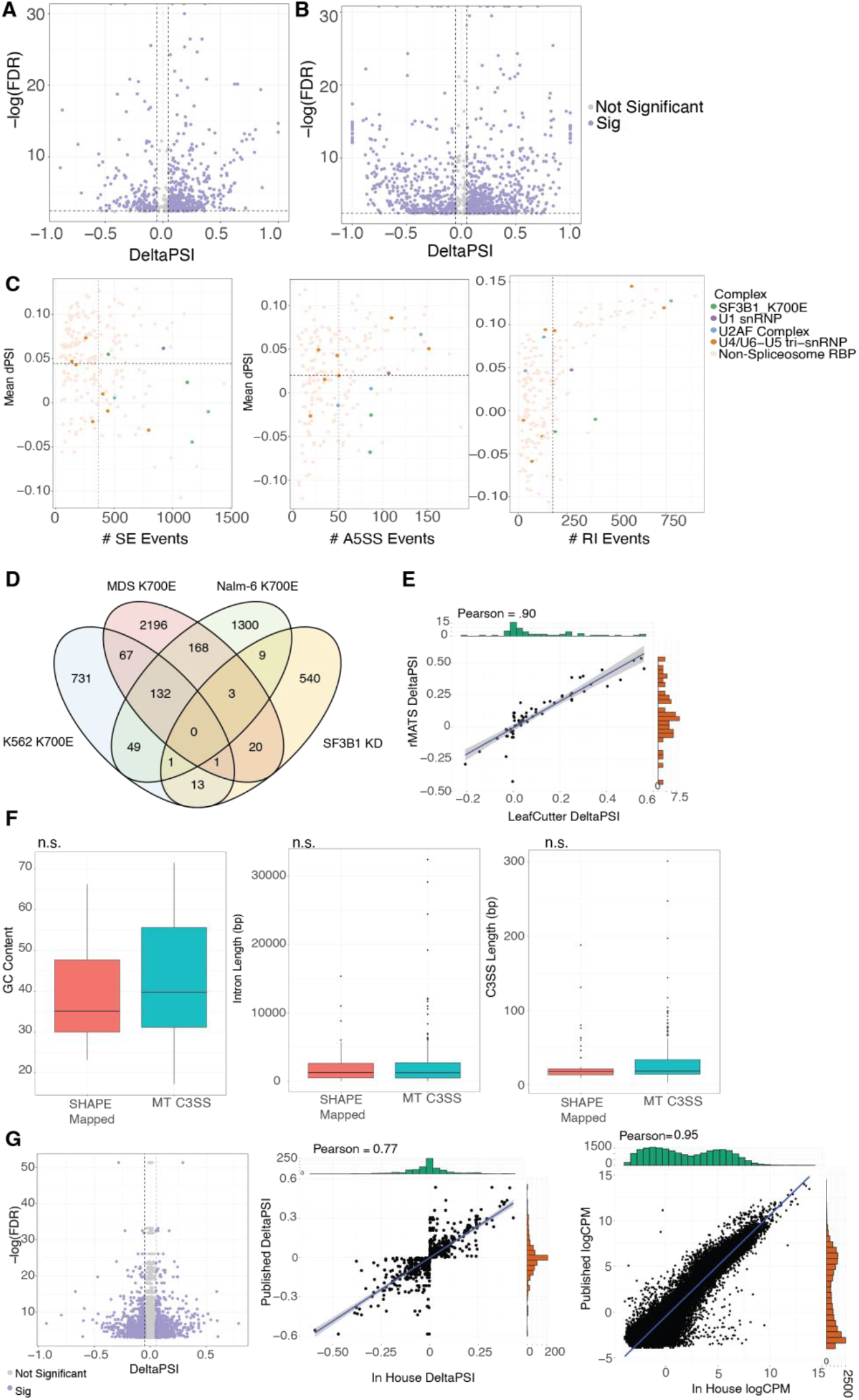
SF3B1 K700E mutants display unique splicing patterns. (A) SF3B1 knockdown in HepG2 (ENCSR896CFV) cells exhibit many alternative splicing events, FDR < .05, deltaPSI > .05. (B) SF3B1 K700E mutation in K562 cells display many alternative splicing events. (C) Little to no trend is observed in exon skipping, alternative 5’ splicing and intron retention events in SF3B1 K700E mutants. Number of and percent spliced in of skipped exons, alternative 5’ splice sites, and intron retention events across ENCODE RBP knockdown series and SF3B1 K700E cell lines. (D) SF3B1 K700E mutants and SF3B1 knockdowns share little overlap in alternative splicing events. Overlap in rMATS C3SS events between SF3B1 K700E cell types and SF3B1 knockdown in HepG2 cells. (E) Alternative splicing analysis with rMATs and LeafCutter display a high correlation. Correlation of deltaPSI in C3SS events called from rMATS and LeafCutter in SF3B1 K700E Nalm-6 cells. (F) Cryptic 3’ splice site sensitive to SF3B1 mutation chosen for structure mapping (n=83) are statistically similar in GC content, intron length, and C3SS length to the entire pool (n=192). (G) In-house sequenced NALM-6 SF3B1 K700E display high correlation in gene expression and alternative splicing profiles with previously published data.

**Supplementary Figure 2.**
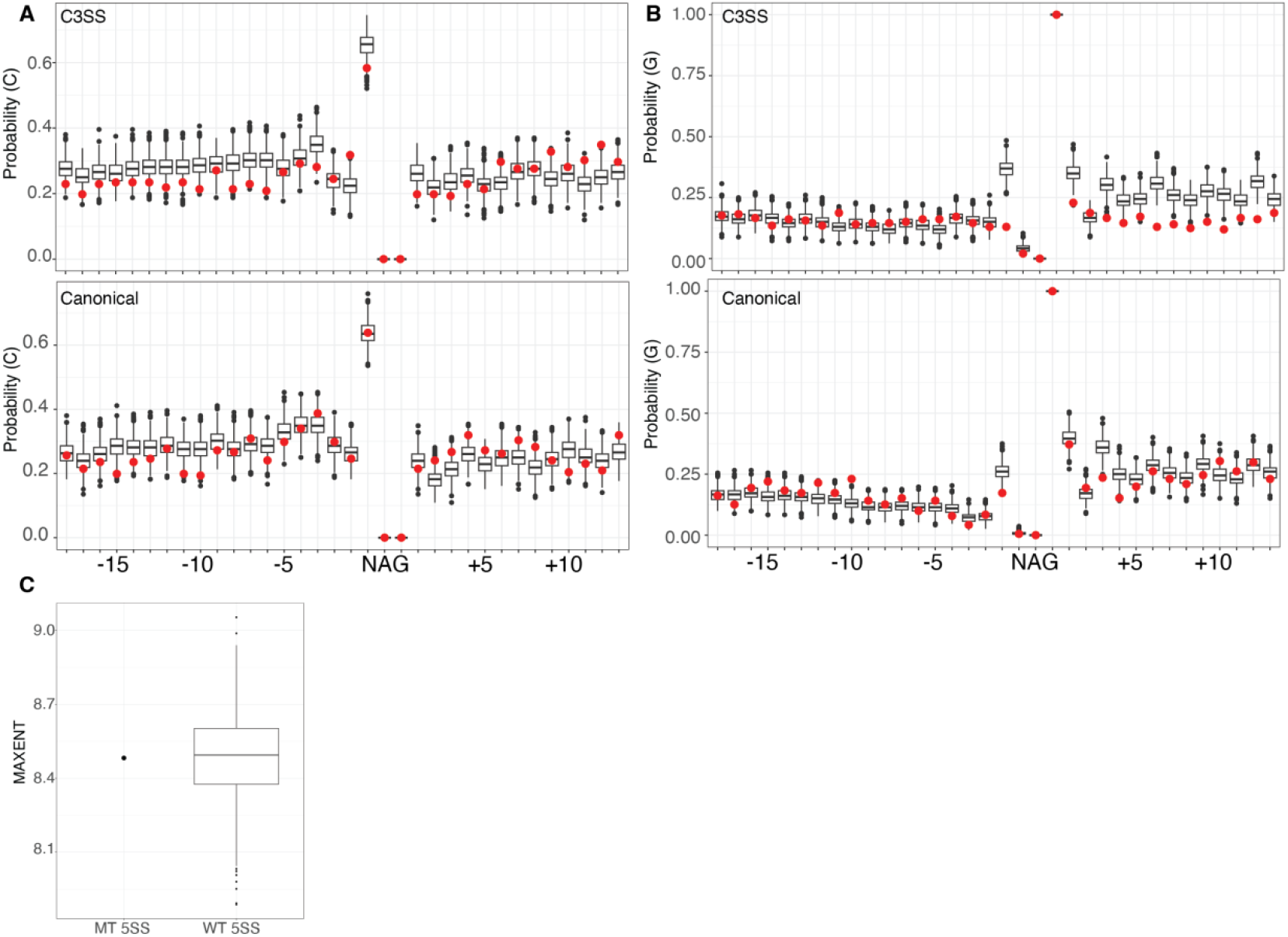
SF3B1 MT C3SS and paired canonical splice sites have similar GC nucleotide composition. (A) SF3B1 MT cryptic 3 splice sites (MT C3SS) and paired canonical display similar probabilities for cytosine and (B) guanine in MT C3SS (top) and paired canonical (bottom) plotted versus randomly bootstrapped WT C3SS and paired canonical. (C) No significant different in the MAXENT of 5’ splice sites paired to SF3B1 MT C3SS and WT C3SS.

**Supplementary Figure 3.**
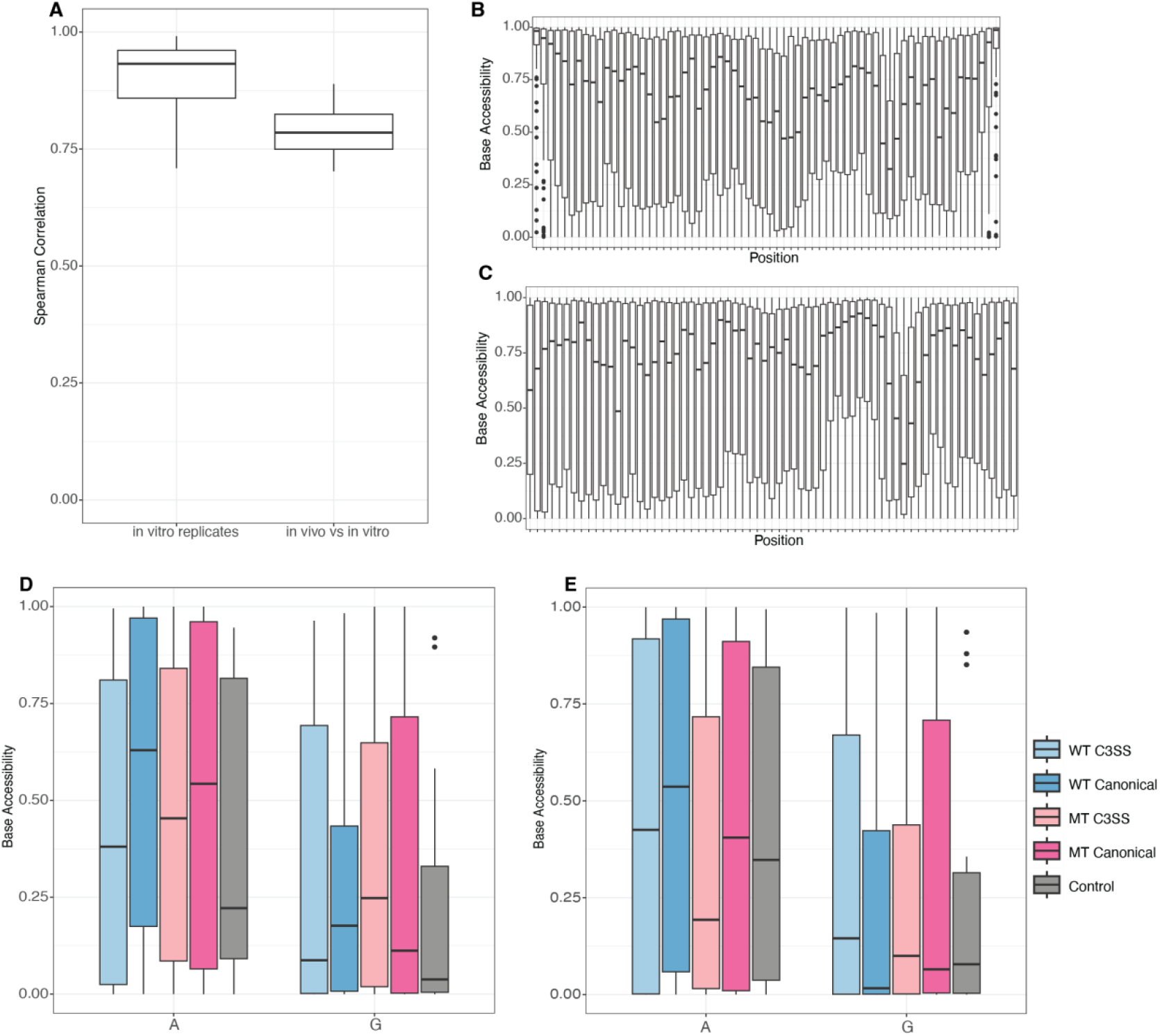
SHAPE correlation and SHAPE guided accessibility patterns. (A) A high spearman correlation is observed between in vitro replicates (n=83) and in vitro versus in vivo structure mapped introns (n=5) > .7. (B) Folding short sequences creates bias in the accessibility of bases at the end of sequences. Base accessibility of MT C3SS of -50 and +15 bases from the splice site. (C) Folding longer sequences while adjusting max distance parameters eliminates end bias observed when folding short sequences. Base accessibility of MT C3SS of - 450 and +450 bases from the splice site with bases -50 and +15 bases from the splice site being visualized. (D) No significant difference in the average base accessibility of cryptic and paired canonical AG splice site motifs folded from -450 and +450 of the splice sites. Base accessibilities were smoothed over max distances of 27 to 34 (D), and 200 to 225 in 5 bp intervals (E), no significant difference between C3SS and paired canonical splice sites were observed.

**Supplementary Figure 4.**
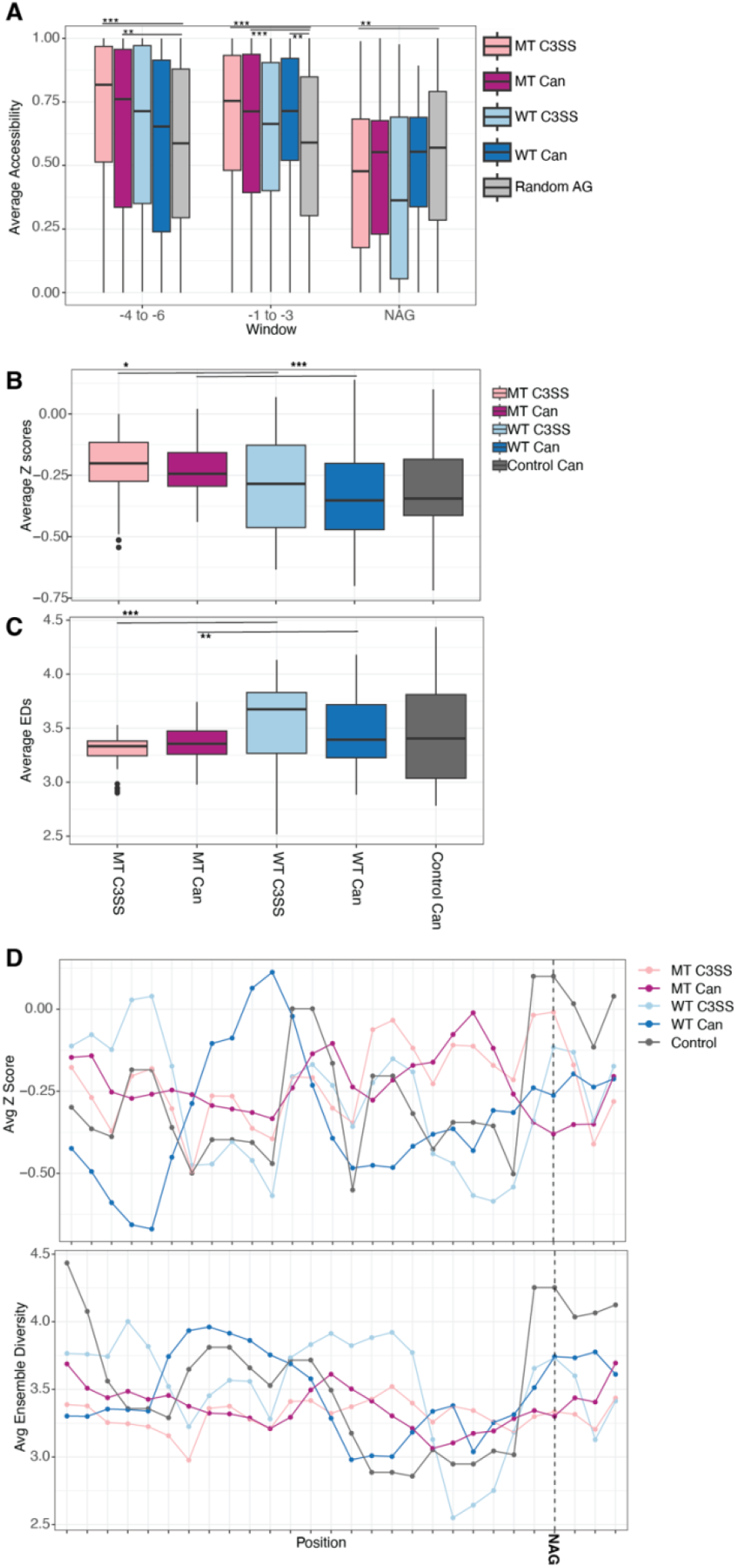
Co-transcriptional model of RNA structure in the proximal intronic regions of 3’ splice sites. (A) SF3B1 MT cryptic 3SS (MT C3SS, light pink) show significantly higher accessibility at -4 to -6, -1 to -3 and at the NAG site compared to random AG dinucleotides (gray). MT paired canonical sites (dark pink) are significant from random AG at -4 to -6 and -1 to -3 windows. Average base accessibility calculated over 3 nt windows up stream of the AG splice site motif. (B) Average Z scores, and (C) average ensemble diversity replicate results from Figure 5C and 5D with max distance of 27 base pairs. Scanfold Z scores and ensemble diversity for - 121 to +15 nt of the AG motifs for categorized splice site types. (D) Z score and ensemble diversity from Scanfold analysis at max distance of 27 nucleotides do not replicate trends seen in Figure 5A and 5B. Average Z scores and average ensemble diversities window averaged three nucleotides at a time for -121 to +15 nucleotides from the AG splice site motif. (*** = p < .0005, **= p < .005, *= p < .05)

